# Age-related differences in fMRI subsequent memory effects are directly linked to local grey matter volume differences

**DOI:** 10.1101/2023.02.23.529668

**Authors:** Jasmin M. Kizilirmak, Joram Soch, Anni Richter, Björn H. Schott

**Author notes:** Correspondence shall be addressed to J.M.K, or B.H.S., /. **Data availability:** The data and analysis scripts/toolboxes used here are available online and linked in the Methods section. Group-level results data are available via OSF.

## Abstract

Episodic memory performance declines with increasing age, and older adults typically show reduced activation of inferior temporo-parietal cortices in functional magnetic resonance imaging (fMRI) studies of episodic memory formation. Given the age-related cortical volume loss, it is conceivable that age-related reduction of memory-related fMRI activity may be partially attributable to reduced grey matter volume (GMV). We performed a voxel-wise multimodal neuroimaging analysis of fMRI correlates of successful memory encoding, using regional GMV as covariate. In a large cohort of healthy adults (106 young, 111 older), older adults showed reduced GMV across the entire neocortex and reduced encoding-related activation of inferior temporal and parieto-occipital cortices compared to young adults. Importantly, these reduced fMRI activations during successful encoding in older adults could in part be attributed to lower regional GMV. Our results highlight the importance of controlling for structural MRI differences in fMRI studies in older adults but also demonstrate that age-related differences in memory-related fMRI activity cannot be attributed to structural variability alone.

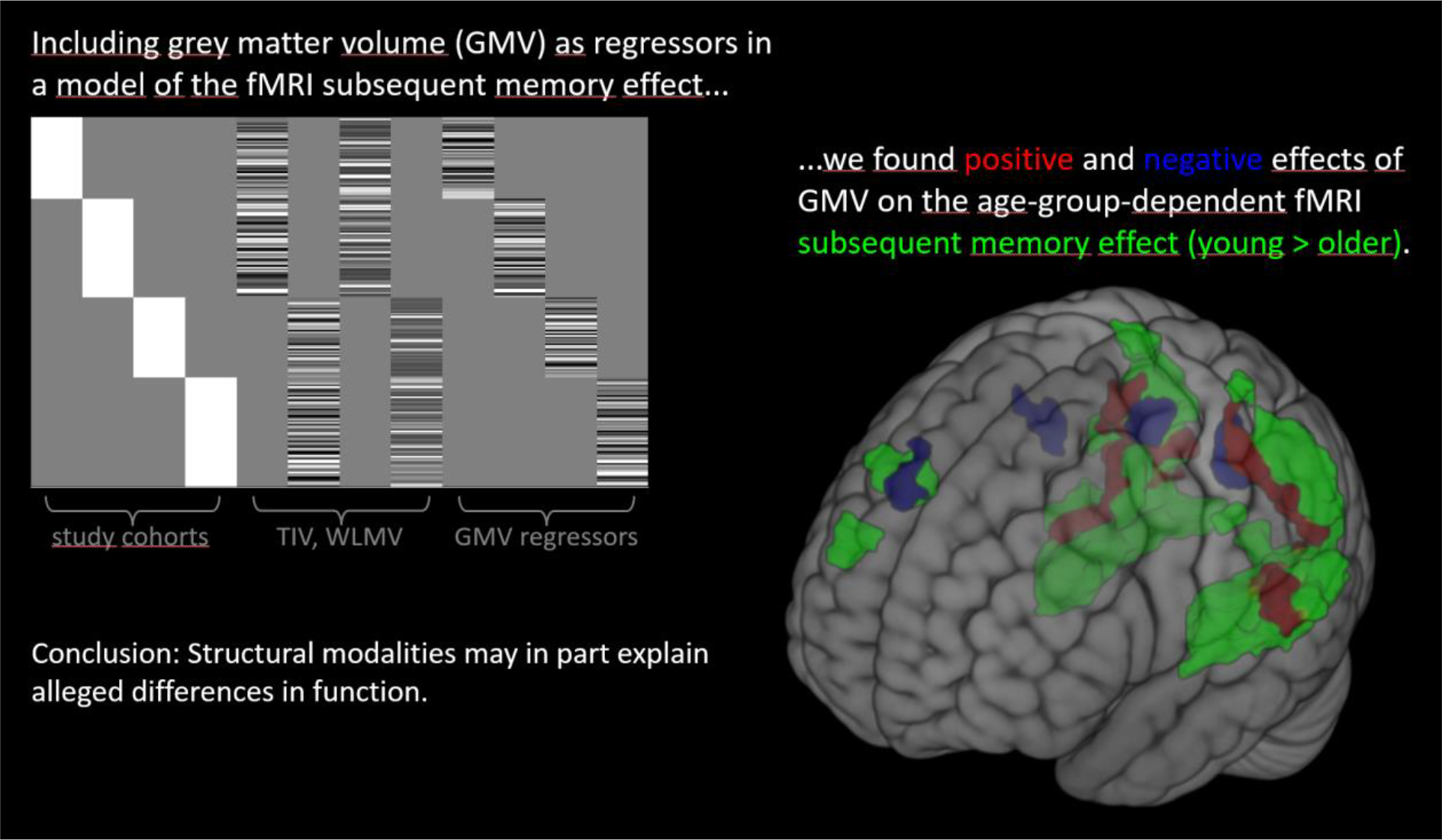

## 1. Introduction

Converging evidence supports an important link between cerebral and cognitive aging, especially with regard to episodic memory function (Cabeza et al., 2005; Nyberg et al., 2012). At an older age, considerable volume loss can be observed. White matter loss is associated with slower reaction times and neurocognitive processing in general, and grey matter loss is associated with a decline in cognitive performance, particularly in attention-demanding, working memory, and episodic memory tasks (Craik and Rose, 2012; Rönnlund et al., 2005). Most studies on the cognitive neuroscience of aging link structural and functional brain changes via behavioral performance measures (Persson et al., 2006), but only few have directly assessed the relationship between local structural and functional age-related differences with multimodal analyses (Boller et al., 2017; Kalpouzos et al., 2012). Direct conclusions on age-related differences in the functional recruitment of brain regions can only be drawn when considering potential links to local differences in brain structure. Here, building on previous analyses of the same data set (Kizilirmak et al., 2023), we assessed the direct, voxel-wise, relationship between functional magnetic resonance imaging (fMRI) correlates of successful episodic memory encoding (subsequent memory effects, SME) and grey matter volume (GMV) in a large sample of young and older healthy adults using multimodal neuroimaging.

## 2. Material and methods

### 2.1. Participants

The sample consisted of 217 neurologically and psychiatrically healthy, right-handed adults who participated after having provided written informed consent. The 106 young participants (47 male, 59 female) had an age range of 18 to 35 years (mean age 24.12 ± 4.00 years). The 111 older participants (46 male, 65 female) had an age range of 60 to 80 years (mean age 67.28 ± 4.65 years). A more detailed description of the study sample has been reported previously (Soch et al., 2021a).

### 2.2. Stimuli, task, and procedure

The paradigm consisted of an encoding and a recognition session with a retention interval of approx. 60 min. Participants incidentally encoded 90 unique photographs of scenes, while performing an indoor/outdoor decision. 88 of the images were presented once during encoding, and two images were familiarized prior to the encoding phase and then repeated 22 times each during encoding, thus allowing for the construction of novelty contrasts (images presented once vs. repeatedly presented ‘master’ images). Recognition memory was tested in a surprise memory test, in which all 90 images from encoding plus 44 new images were presented. Participants provided recognition-confidence ratings (5-point rating scale: ‘sure new’ over ‘undecided’ to ‘sure old’). For a more detailed description, see (Soch et al., 2021a).

### 2.3. MRI data acquisition

Structural and functional MRI data was acquired on two Siemens 3T MR scanners (Siemens Verio: 58 young, 64 older; Siemens Skyra: 48 young, 47 older). A T1-weighted magnetization-prepared rapid gradient echo (MPRAGE) image (TR = 2.5 s, TE = 4.37 ms, flip-α = 7°; 192 slices, 256 x 256 in-plane resolution, voxel size = 1 x 1 x 1 mm) was acquired for regional grey matter assessment via voxel-based morphometry (VBM) and for optimized normalization of fMRI data. Functional data comprised T2*-weighted echo-planar images (EPI; TR = 2.58 s, TE = 30 ms, flip-α = 80°; 47 axial slices, 64 x 64 in-plane resolution, voxel size = 3.5 x 3.5 x 3.5 mm) which were acquired while participants performed the encoding phase of the subsequent memory paradigm (8:51 min; 206 EPIs). Additionally, phase and magnitude fieldmap images were acquired to improve correction for artifacts resulting from magnetic field inhomogeneities. T2-weighted FLAIR images (192 sagittal slices, TR = 5.0 s, TE = 395 ms, 256 x 256 mm in-plane resolution, voxel size = 1.0 x 1.0 x 1.0 mm) were acquired to assess white matter lesions.

### 2.4. MRI data analysis

VBM was used for the assessment of voxel-wise GMV and total intracranial volume (TIV), as implemented in the Computational Anatomy Toolbox (CAT12, Gaser and Dahnke, 2016), following a previously described protocol (Assmann et al., 2021). Segmented GM maps were normalized to the Montreal Neurological Institute (MNI) reference frame, using the SPM12 DARTEL template, employing a Jacobian modulation, and keeping the spatial resolution 1 mm isotropic. Normalized GM maps were smoothed with a Gaussian kernel of 6 mm at FWHM. FLAIR images were used to assess white matter lesion volume (WMLV) via automatic segmentation using the Lesion Prediction Algorithm (Schmidt, 2017). Assessing WMLV was important to control for age-related differences in fMRI due to WM lesions.

Preprocessing of fMRI data was carried out using Statistical Parametric Mapping (SPM12; Wellcome Trust Center for Neuroimaging, University College London, UK). EPIs were corrected for acquisition time delay (*slice timing*), head motion (*realignment*) and magnetic field inhomogeneities (*unwarping*), using voxel-displacement maps derived from the fieldmaps. The MPRAGE image was co-registered to the mean unwarped image and segmented into six tissue types, using the unified segmentation and normalization algorithm implemented in SPM12. The resulting forward deformation parameters were used to normalize unwarped EPIs into a standard stereotactic reference frame (MNI; voxel size = 3 x 3 x 3 mm). Normalized EPIs were spatially smoothed using a 6 mm isotropic Gaussian kernel.

To assess the fMRI subsequent memory effect, we specified a general linear model (GLM) for the encoding-related fMRI data. Two onset regressors for unique images (“novelty regressor”) and familiarized images (“master regressor”) were created as box-car stimulus functions (2.5 s), convolved with the canonical hemodynamic response function. The novelty regressor was parametrically modulated with the arcsine-transformed behavioral response rating from the recognition memory test, yielding a parametric subsequent memory regressor (Soch et al., 2021b). The model further included the six rigid-body movement parameters obtained from realignment as covariates of no interest, and a constant representing the implicit baseline. A positive effect of the parametric modulator reflects differential activation for remembered vs. forgotten items and is referred to as the subsequent memory effect (SME).

### 2.5. Multi-modal MRI data analysis

Multi-modal analyses were conducted using a toolbox^1^ developed for use with SPM12. First-level fMRI contrast maps reflecting the parametric SME were used as the dependent variable, to be explained by two categorical factors *age group* (young, older) and *scanner* (1=Verio, 2=Skyra). Scanner was included to account for potential effects of different head coils, though none were found. Continuous covariates of no interest were TIV and WMLV, separately modeled for each age group. Importantly, we further included one GMV regressor per age group^2^ x scanner combination to add GMV as an explanatory variable. GMV maps were resampled to match the resolution of the functional images. The design matrix is shown in Figure 1D. The model represents an adaptation of model 2 from Kizilirmak et al. (2023):

**Figure 1.**
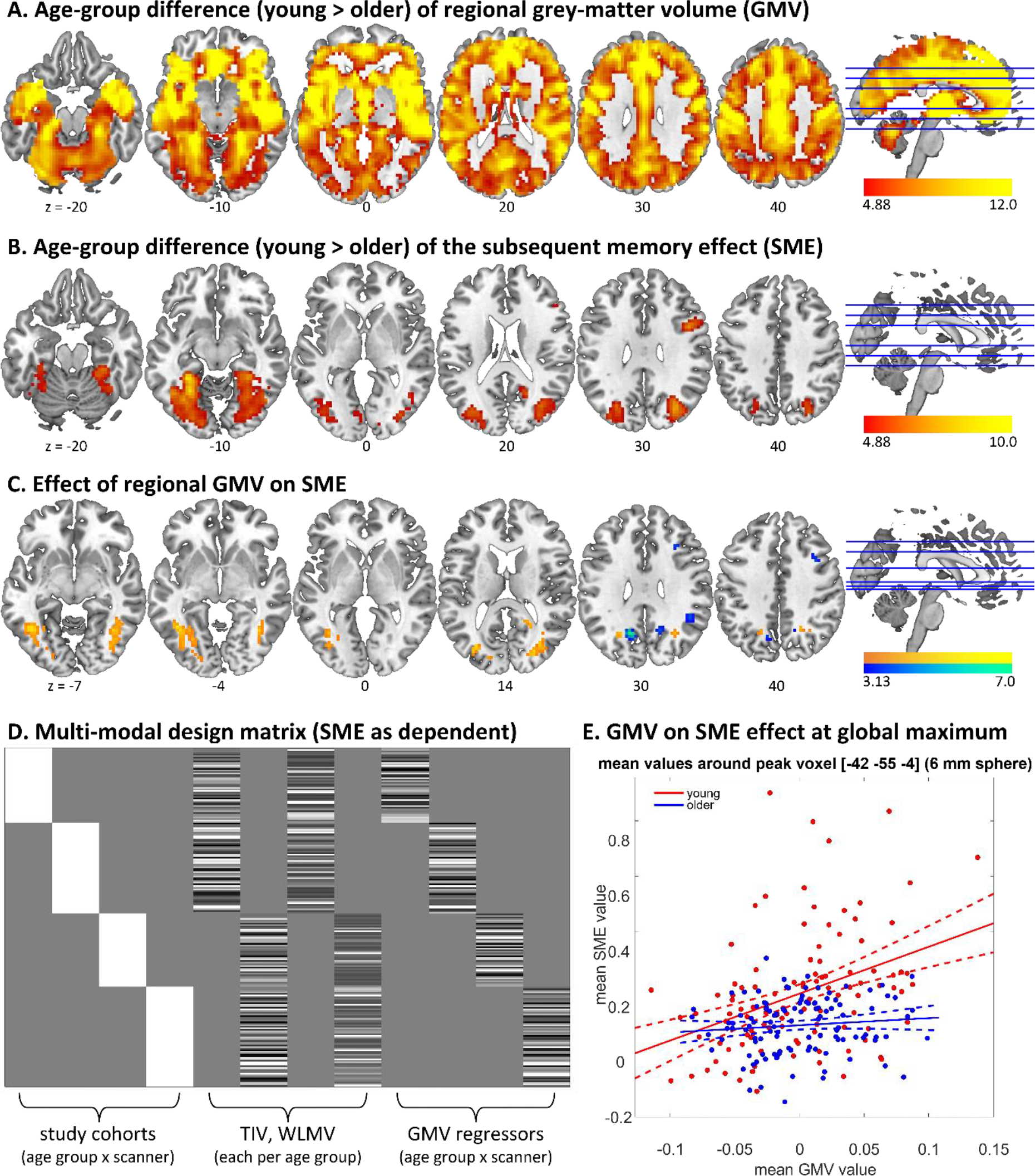
Effects of age differences in grey matter volume (GMV) and the fMRI subsequent memory effect (SME) as well as influences of GMV on SME. A. GMV contrast young > older. B. SME contrast young > older. C. Positive (warm colors) and negative (cold colors) effect of GMV on SME. D. Design matrix for the model used in B and C. Panels A and B show voxel-level FWE-corrected effects at p < .05, cluster threshold 10 voxels, panel C shows effects at cluster-level FWE-corrected p < .05. E. The positive effect of GMV on SME plotted for the mean from a 6 mm sphere around the global maximum [-42 −55 −4].

SME ∼ age_group(young, old) + scanner(1, 2) + TIV + WMLV + GMV.

All MATLAB scripts for the GLMs are provided at OSF (https://osf.io/cxm2f/) and MRI data (GMV maps and SME contrast maps) at NeuroVault^3^. The significance level was set to p < .05, corrected for family-wise error (FWE) at cluster level, cluster-defining threshold of p < .001, uncorrected. A combined binarized active voxel mask for SME and GMV was used.

## 3. Results

First, we assessed age-group differences in GMV and SME separately. As expected, there were pronounced differences in both cortical and subcortical GMV across the whole brain in the young > older contrast (Figure 1A), while few differences were found in the reverse contrast (young < older), mainly along the GM-WM border. As reported previously (Kizilirmak et al., 2023; Soch et al., 2021a), the age differences in SME included regions along the ventral and dorsal visual stream showing for young > older adults (Figure 1B) and mostly regions of the default mode network for young < older adults, whereby the first differences are due to higher activations in younger adults and the latter due to lower deactivations in the older adults. We then assessed the contribution of GMV to the observed SME (Table 1). Our multi-modal analysis revealed positive as well as negative effects of GMV on SME (Figure 1C). Notably, the positive effects, i.e., higher GMV associated with a higher SME, were observed in overlapping occipital and lingual gyrus regions as the age effects on the SME itself (cf. Figure 1B vs. 1C). Negative effects, i.e., higher GMV associated with lower SME, were especially found in bilateral precuneus.

**Table 1.**
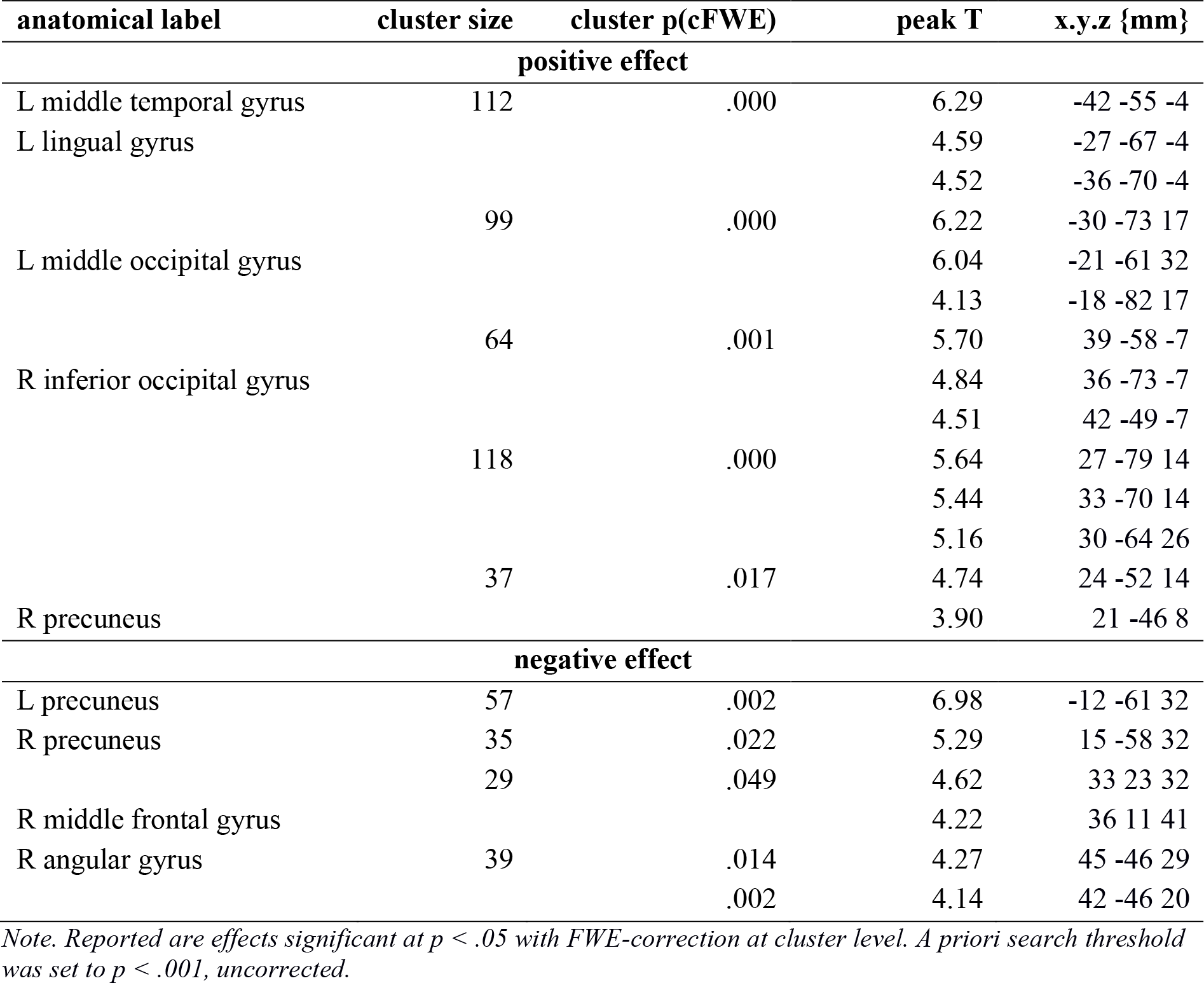
Positive and negative effects of GMV on the fMRI subsequent memory effect.

As it may be another source for functional age differences due age-dependent structural changes, we lastly assessed potential effects of WMLV. However, there were none, not even at a lowered significance threshold of p < .001 unc. and a cluster threshold of 10 voxels.

## 4. Discussion

Here, we directly assessed the relationship between local structural (GMV) and functional age-related differences during SME. To our knowledge, there has only been one study thus far with a similar methodological approach (Kalpouzos et al., 2012). In 16 young (8 female) and 20 older healthy participants (all female), the authors investigated the influence of voxel-wise GMV on intentional episodic memory encoding. They masked the fMRI contrast maps with the thresholded encoding-versus-control contrast and assessed the main effect of GMV. They, too, found that occipital GMV loss accounted for an underrecruitment during encoding, but could only infer this indirectly, as they were unable to differentiate positive and negative effects due to restrictions of the toolbox used (BPM; Casanova et al., 2007). In the present study, we followed up on this valuable multimodal approach using data from a larger and more representative healthy population (106 young and 111 older adult, balanced gender distribution in both age groups), assessing both positive and negative effects of GMV on fMRI encoding contrasts. Further, fMRI activity related to encoding was assessed as a parametrically modelled subsequent memory effect, which reflects *successful* encoding and is more specific than an encoding-versus-control contrast.

Older adults showed reduced activations during successful encoding of visual scenes in the same regions in which they show significant reductions in GMV, that is, occipital and lingual gyrus, as well as middle temporal lobe. Moreover, we found a direct association between regional GMV and successful encoding-related fMRI. GMV can be indicative of the metabolic capacity of brain regions. Regions with lower gray matter may have lower energy and oxygen demands, possibly due to fewer neurons and synapses, which would in turn likely lead to lower levels of neural activity. Moreover, older adults often show quantitatively, and, to some extent, even qualitatively distinct patterns of brain activity compared to younger adults, potentially compensating for age-related structural changes (see Maillet and Rajah, 2014, for a meta-analysis). Notably, the GMV effects on the fMRI subsequent memory effect were mostly found in posterior brain regions despite the GMV age differences being most pronounced frontally (Figure 1A). This may be explained by the use of a visual encoding task and is in line with a recent study, in which angular gyrus GMV was found to predict differences in memory precision in older adults (Korkki et al., 2023). An inverse relationship between GMV and SMV was found in the precuneus and medial PFC (Table 1). This may be best explained by the fact that these regions show negative subsequent memory effects (Kizilirmak et al., 2023), such that the correlation with absolute BOLD amplitudes would be positive in these regions. Other studies failed to find an association between GMV and fMRI contrasts (Pur et al., 2019), which may be due to the use of ROI-wise GMV means, pointing out the importance of assessing the regional variation in a more fine-grained, e.g., voxel-wise manner. However, Pur et al. also found an association between structural and functional MRI in terms of cortical thickness predicting BOLD standard deviation in a reaction time task.

On this basis, our results speak for including structural modalities in the models when comparing functional MRI effects of participant groups with differences in brain structure to detect potential confounds that may in part explain alleged differences in function. Lastly, however, age-related differences in fMRI subsequent memory effects are robust, even when individual regional GMV differences are accounted for (Figure 1B). This is assuring regarding the many studies on age differences in episodic memory formation, but cannot by itself be generalized to other participant groups, stimulus modalities and fMRI contrasts.

## Acknowledgements

This study was supported by the State of Saxony-Anhalt and the European Union (Research Alliance “Autonomy in Old Age” to A.R. and B.H.S.) and by the Deutsche Forschungsgemeinschaft (SFB 1436, TP A05). We thank Hartmut Schütze for programming the experiment, Hannah Feldhoff, Larissa Fischer, Lea Knopf, Matthias Raschick, and Annika Schult for helping with participant recruitment and data collection, and Kerstin Möhring, Katja Neumann, Ilona Wiedenhöft, and Claus Tempelmann for help with MRI acquisition.

## Footnotes

1 https://github.com/JoramSoch/MMA

2 Effects of GMV on SME per age group are not reported here due to space limitations, but thresholded contrast images can be found in our OSF repository.

3 https://neurovault.org/collections/XDYPPLTD/

